# Musical Imagery Involves the Wernicke’s Area in Bilateral and Anti-Correlated Network Interactions in Musicians

**DOI:** 10.1101/142950

**Authors:** Yizhen Zhang, Gang Chen, Haiguang Wen, Kun-Han Lu, Zhongming Liu

**Author notes:** Correspondence Zhongming Liu, PhD Assistant Professor of Biomedical Engineering, Assistant Professor of Electrical and Computer Engineering, College of Engineering, Purdue University, 206 S. Martin Jischke Dr., West Lafayette, IN 47907, USA.

## Abstract

Musical imagery is a human experience of imagining music without actually hearing it. The neural basis of such a mental ability is unclear, especially for musicians capable of accurate and vivid musical imagery due to their musical training. Here, we visualized an 8-min symphony as a silent movie, and used it as real-time cues for musicians to continuously imagine the music for multiple synchronized sessions during functional magnetic resonance imaging. The activations and networks evoked by musical imagery were compared with those when the subjects directly listened to the same music. The musical imagery and perception shared similar responses at bilateral secondary auditory areas and Wernicke’s area for encoding the musical feature. But the Wernicke’s area was involved in highly distinct network interactions during musical imagery vs. perception. The former involved positive correlations with a subset of the auditory network and the attention network, but negative correlations with the default mode network; the latter was confined to the intrinsic auditory network in the resting state. Our results highlight the important role of the Wernicke’s area in forming vivid musical imagery through bilateral and anti-correlated network interactions, challenging the conventional view of segregated and lateralized processing of music vs. language.

## Introduction

Mental imagery often occurs as experiences such as “seeing in the mind's eye”, or “hearing in the head”. It refers to quasi-perceptual mentation that resembles vivid perception in the absence of any external stimuli ^1^. Questions remain largely unanswered as to how the brain generates imagery, and whether imagery and perception share a similar or distinct mechanism ^2–4^. In this regard, previous studies have been focused more on the visual ^5–9^ and motor systems ^10–13^, rather than the auditory system ^14,15^. However, unlike visual or motor imagery, auditory imagery may restore real sensation in mind with surprisingly high accuracy ^16–20^. As a well-known demonstration, Ludwig van Beethoven, who was deaf in his later years, went on composing and conducting great symphonies guided by his “inner ear” ^21^. Although nearly everyone has such experiences as “singing a song in mind”, musicians excel in such mental ability through musical training, e.g. being able to have an orchestra in mind to perform a mental concert while reading a music sheet ^22^.

Previous studies have shown that musical imagery preserves many auditory and musical features ^16–18^. Initial evidence supports the notion that musical imagery and perception may share a similar neural basis within and beyond the auditory system ^23–25^. During both musical perception and imagery, cortical activations are observable at the secondary auditory cortex, the association cortex ^17,20,24,25^, and even the motor cortex especially for musicians ^14,26–28^. In addition, musical imagery in musicians is of interest for studying experience-dependent neural plasticity, since musical training reshapes neuronal networks to promote auditory imagery ^29,30^. Thus, imaging and understanding musical imagery in musicians holds a unique value for sensory and cognitive neuroscience.

In this study, we aimed to map and compare cortical networks during musical perception and imagery using functional magnetic resonance imaging (fMRI). Uniquely, we used music visualization to guide continuous and complex musical imagery with well-controlled timing, unlike prior studies that focus on short and simple imagery tasks ^15,20,24^. This musical-imagery task allowed us to map task-activated regions and networks based on model-free analyses of intra-subject reproducibility and functional connectivity ^31,32^. Task-evoked activity was also compared with musical feature time series, to reveal the music information preserved during imagery as well as during perception. Lastly, the task-activated networks were further compared to intrinsic functional networks in the resting state. Taken together, our unique study design and advanced analyses aimed to shed mechanistic insights onto the neural-network basis of musical imagery.

## Results

### Sustained musical imagery evoked wide-spread cortical activation

When a subject imagined a sustained music piece while watching a movie that visualized the music, the visual input served to control the timing and inform the content of musical imagery (Fig. 1a). With such real-time visual cues, the subject was able to consistently imagine the same 8-min music piece for 12 repeated sessions during fMRI scans. In this experimental design, the cortical activation during sustained and complex musical imagery was mapped by assessing the reproducibility in the fMRI signal across different sessions of the same imagery task. Fig. 1c shows the cortical activation with significant intra-subject reproducibility measured as the inter-session correlation. The activated areas covered a large part of the cortex, including the primary visual cortex, dorsal-stream visual areas, the attention network in the association cortex, the secondary auditory cortex, and the Wernicke’s area (Fig. 1c). These areas were likely activated by different sensory and cognitive processes given the visually-cued musical imagery involved both visual perception and auditory imagery.

**Figure1.**
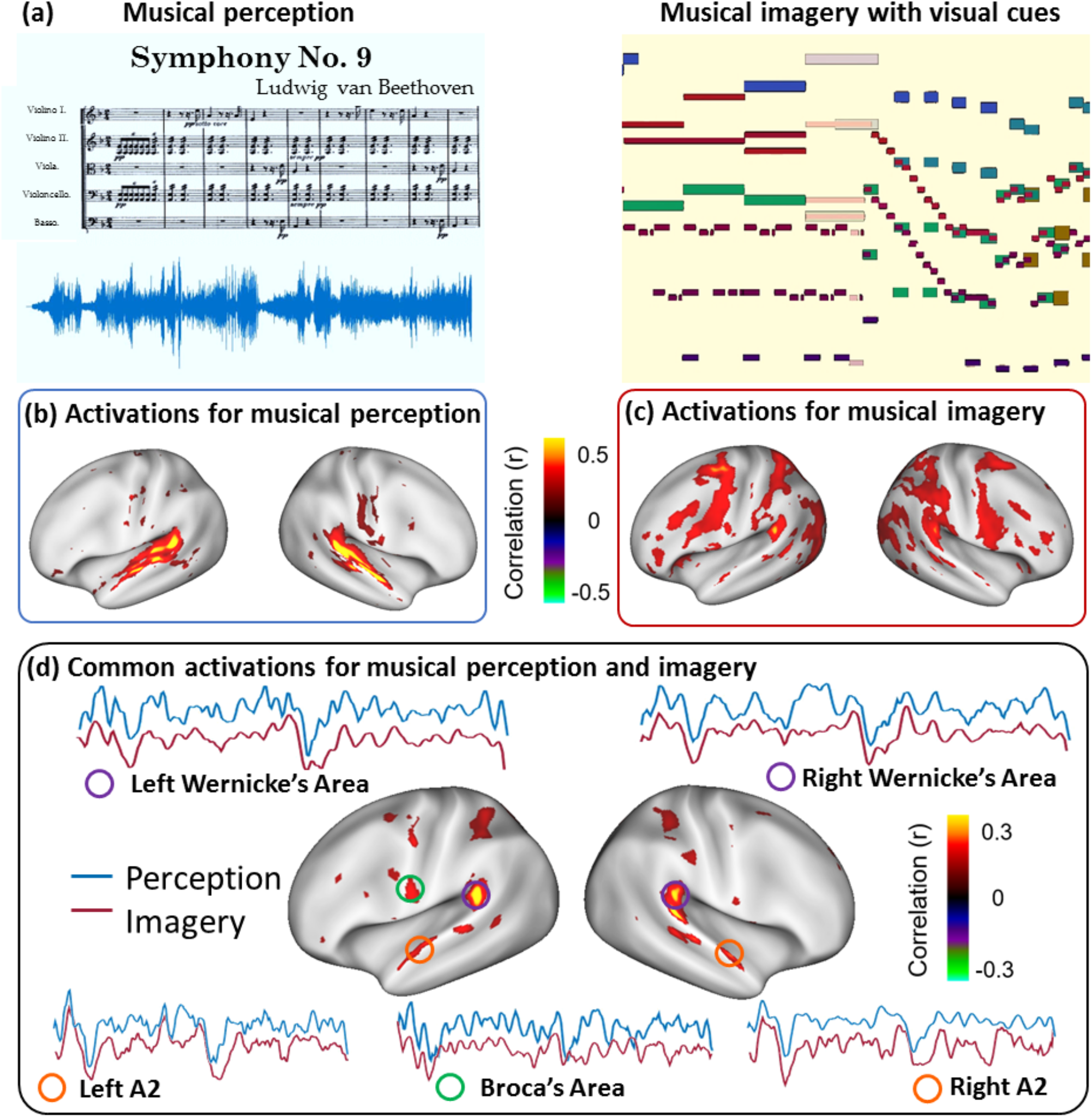
Distinct and common cortical activations with musical perception and imagery. (a) paradigm for musical perception (left) and imagery (right). The visualized music is an animation with bars moving from right to left as the music flows. It includes all the musical information as in a standard music sheet: the length of the bars indicates the note length (rhythm and duration); the height of the bars indicates the keynote (pitch); the color of the bars indicates the instrument (timbre). (b) cortical activations for musical perception (two-tailed significance level p<0.01). (c) cortical activations for musical imagery (two-tailed significance level p<0.005). (d) shared common cortical substrates between musical perception and musical imagery (two-tailed significance level p<0.01). The time series was extracted from the fMRI signal of the averaged perception and imagery sessions from labeled locations.

### Musical imagery and perception shared common cortical substrates

Our next step was to dissociate the imagery-related activations from those due to visual perception. For this purpose, the same music piece was directly presented to the subject in the absence of any visual input. With the musical perception, the intra-subject reproducibility of the fMRI signal revealed cortical activations in a much smaller extent mostly confined to the auditory system (Fig. 1b). The auditory-evoked activations were most notable in the primary and secondary auditory cortex, as well as the Wernicke’s area (Fig. 1b). The fact that the secondary auditory cortex and the Wernicke’s area were activated by both musical imagery and perception, suggest that they might be the common cortical substrates for both auditory perception and imagery.

To test this hypothesis, we assessed for each voxel the correlation in voxel time series between every musical-imagery session and every music-perception session. This analysis directly revealed the areas that showed consistent responses to both the imagery and perception tasks (Fig. 1d). Such areas included the bilateral Wernicke’s areas and secondary auditory cortex (or belt areas), to a less degree the association cortex. At each of the co-activated areas, the response time series over an 8-min period was seemingly irregular, but highly similar whether the musical content was delivered in an auditory or visual form (Fig. 1d). Similar phenomena were observed for all three subjects (Fig. 2).

**Figure2.**
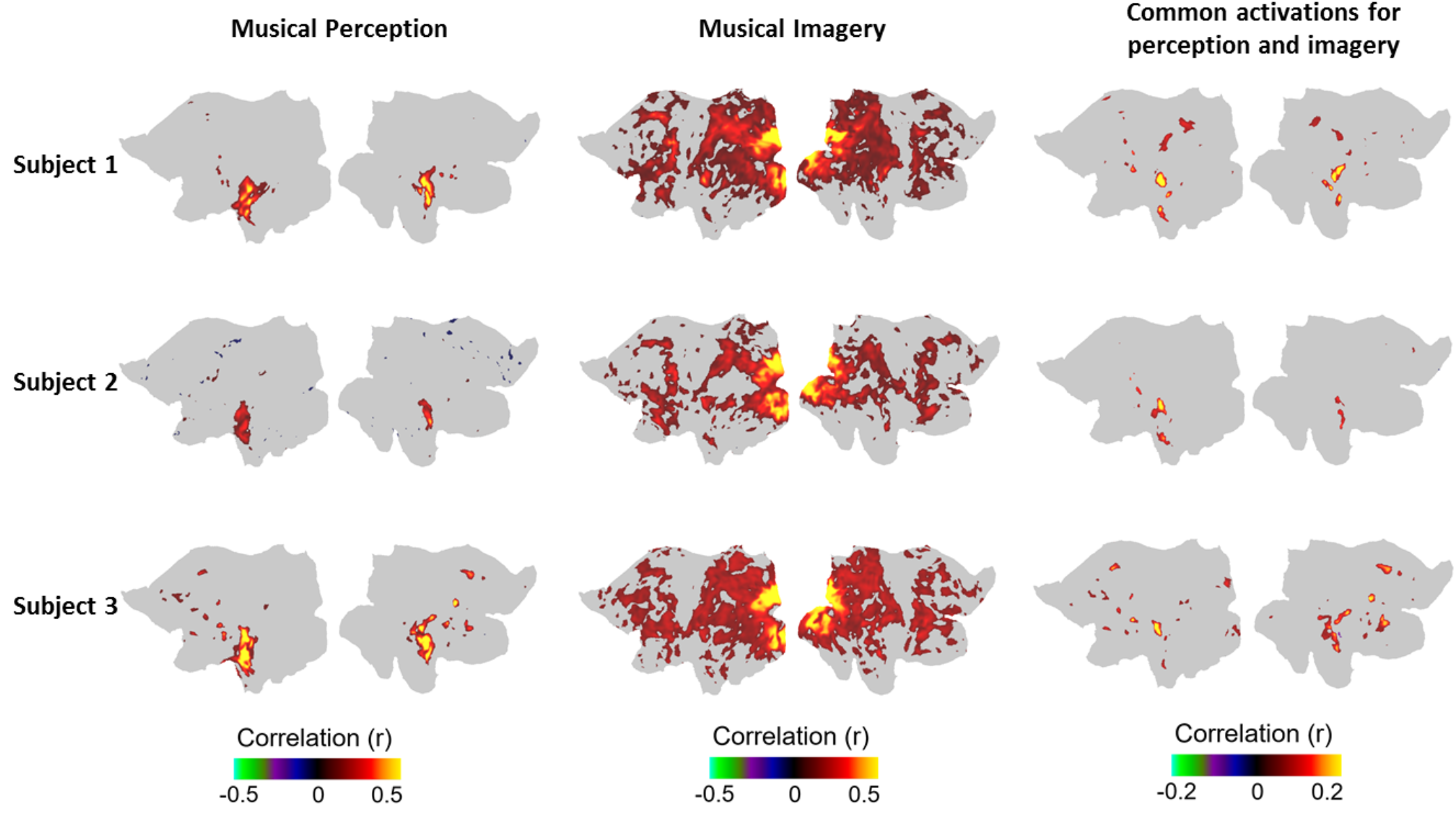
Cortical activations for musical perception and imagery was consistent across all subjects. Each row indicates the cortical mapping of one subject. The first column is the cortical activations for musical perception; the second column is the cortical activations for musical imagery; the third column is the shared common cortical substrates between the two tasks. The first row (subject 1) is a flattened view of surface mapping as Fig1. (b) (c) (d).

### Responses at Wernicke’s areas coded musical features during imagery

We further asked what musical information was preserved in the brain during musical imagery. To address this question, we explored a specific auditory feature known as the spectral flux. This feature reflects how quickly the power spectrum of a sound wave changes over time ^33^. As shown in Fig. 3.a, extracting the spectral flux from the music stimulus resulted in a feature time series that represented the musical information. After convolving it with the HRF, the musical-feature time series was correlated to the fMRI signal at every voxel during musical imagery. Consistent across subjects, the significantly correlated areas included early visual areas and the Wernicke’s areas in both hemispheres (Fig. 3b, right). The former was perhaps not surprising due to the real-time visual cues that changed along with the music information. The latter was non-trivial and interesting, in particular because the responses at the same Wernicke’s areas were correlated with the spectral flux feature when the music was directly delivered to and heard by the subjects (Fig. 3.b, left). The Wernicke’s areas were consistently correlated with the musical-feature time series for both imagery and perception and across all subjects.

**Figure3.**
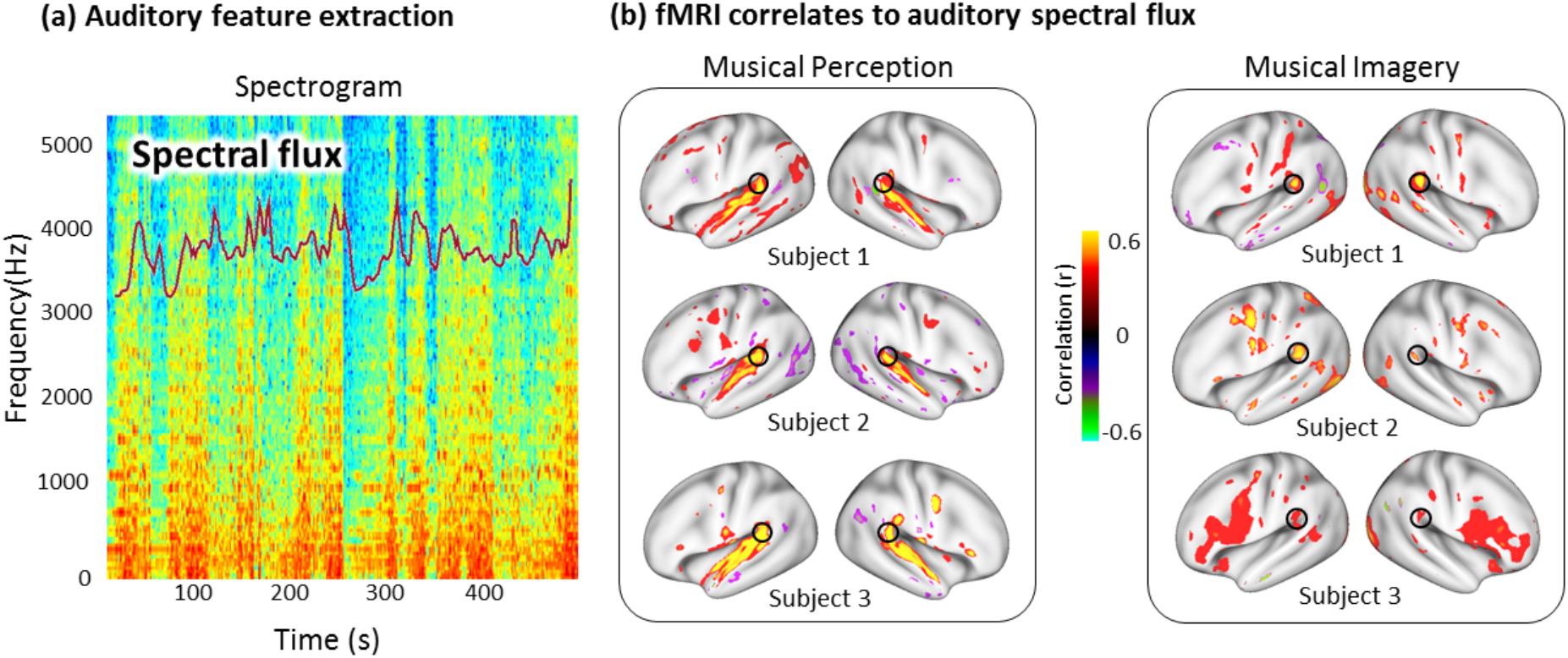
Responses at Wernicke’s areas coded musical features during imagery. (a) The auditory spectral flux was extracted from the stimulus spectrogram as a feature showing how quickly the power spectrum of a sound wave changes over time. (b) Spectral flux was highly correlated with the fMRI signals in the common cortical regions shared between musical perception and imagery, especially in early visual areas and the bilateral Wernicke’s areas (as circled on the maps).

### Musical imagery and perception evoked highly distinct cortical networks

We further asked how the brain engaged, similarly or differently, its functional networks to support musical imagery and perception. To answer this question, we mapped and compared the task-evoked patterns of functional connectivity during the musical imagery versus perception task. This analysis was based on seed-based inter-session functional connectivity attributable to task-evoked responses instead of ongoing activity during the task ^34^. Four seed locations were chosen from the bilateral Wernicke’s areas and secondary auditory cortex, because these areas were activated by both the musical imagery and perception tasks, as shown in Figs. 1 & 2. Surprisingly, the seed-based patterns of task-evoked functional connectivity were highly distinct between the musical perception task and the musical imagery task (Fig. 4). In contrast to a unimodal network involved in musical perception (Fig. 4, left), musical imagery recruited broader activations from the attention network in the association cortex, Wernicke’s areas, along with deactivations of the default mode network (Fig. 4, middle). The difference was notable for each of the four seed locations. The only common pattern of functional connectivity shared between musical imagery and perception was the strong mutual correlation between the Wernicke’s areas and the secondary auditory cortex for both of the tasks. In comparison with the resting state networks, the networks evoked by musical perception agreed with the corresponding intrinsic functional networks (Fig. 4, right). The networks evoked by musical imagery were not confined to any single intrinsic network; instead they all involved complex interactions among multiple networks, including higher-order auditory networks, the attention network, and the default-mode network.

**Figure4.**
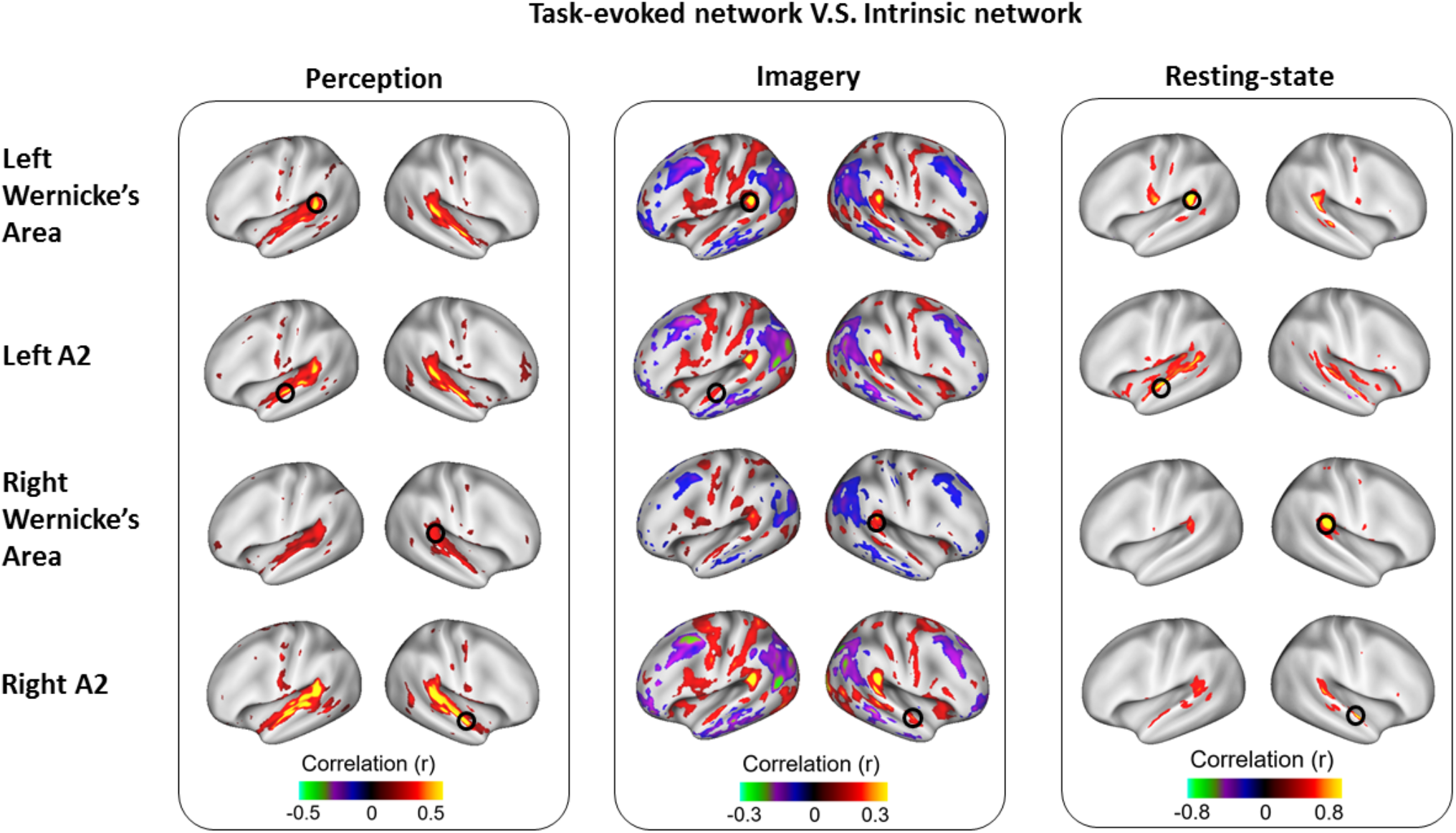
Task-evoked cortical network versus intrinsic network. The perception-(first column) and imagery-evoked (second-column) cortical networks were mapped by cross-session seed-based correlation in the task-related fMRI signals. The third column is the intrinsic network mapped from resting-state fMRI. Each row indicates one specific seed location (circled on the maps), labelled on the left.

## Discussion

In this study, we introduce a new stimulus paradigm to map and compare cortical activations and networks in musicians during sustained and complex musical imagery vs. perception. During visually-cued musical imagery, musical information is transferred from its visual form via the activation of the attention network and the deactivation of the default-mode network. The musical information is conveyed to and coded by bilateral Wernicke’s areas and the secondary auditory areas, highlighting their critical roles in forming vivid musical imagery resembling the experience of listening to music. However, these areas engage in highly distinct network interactions to support musical imagery vs. perception: being much more widespread, complex, and multi-modal for musical imagery, rather than being confined to intrinsic and unimodal networks for musical perception.

Imagery is much harder to control than perception. The experience of imagery is subjective, isolated, and difficult to measure, especially for sustained and complex imagery. This challenge limits previous studies to have only used simple and short auditory imagery tasks in a block or event-related design ^20,35^, and/or subjects’ cognitive self-assessment of their compliance to the imagery task ^20,24^. These limitations hinder the repeated measures of cortical activity underlying any complex imagery such as listening to a symphony in mind for a long period. In turn, the lack of repeated measures limits the ability to study regional and network-level dynamics and coding that supports vivid musical imagery.

Unique to this study is the use of music visualization to control the timing and inform the content of complex and sustained musical imagery. The visualization uses a similar concept as Karaoke, enabling subjects to repeatedly imagine the same music with a controlled tempo. In this way, mental activity is synchronized across repeated imagery experiments. It allows the fMRI scans to be compared across repeated sessions for mapping the imagery-evoked activations and networks in a similar fashion as cortical mapping during natural vision or hearing^31^. Moreover, this stimulus paradigm also allows the synchronization between imagining the music and listening to the music.

Thus, imagery-evoked activity and perception-evoked activity can be directly compared to map the cortical representations shared between musical imagery and perception.

Beyond mapping imagery-evoked activations and networks, this visually-cued imagery paradigm also offers a new opportunity to study neural coding or develop mind-reading for imagery. In particular, brain decoding methods require extensive data for training and testing decoding models^36,37^. Such data are often unavailable, since the content of imagery is either inaccessible or inaccurate. Being able to control the imagery content with temporally synchronized brain activity measurements is desirable to generate data for rigorously training and testing decoding models, before attempting to apply them to more complex imagery or cognitive processes, such as dreams^9^.

Our results suggest that two areas are essential to musical imagery: secondary auditory cortex and Wernicke’s area. The involvement of the secondary auditory cortex in musical imagery echoes and extends the prior findings ^14,17,20^. However, it was unexpected that the Wernicke’s area on the left hemisphere and its homologous area in the right hemisphere were both activated with similar responses during musical imagery and perception (Fig. 1). Conventionally, the Wernicke’s area is thought to be responsible for speech or language processing^38,39^. Music and language involve largely different cortical areas and processes ^40^. However, these conventional views have been challenged, as findings from recent imaging and neurophysiological studies shed light on the common neural basis shared between language and music ^41^. Indeed, our results suggest that the Wernicke’s area plays an important role in forming vivid musical imagery resembling the experience of listening to music.

The engagement of the Wernicke’s area in musical imagery might be unique to musicians. Note that the stimulus in this study is a piece of classical music without lyrics. It is thus unlikely that the response at the Wernicke’s area reflects language comprehension. However, for musicians “reading the visualized music” or “hearing the music” might involve similar cognitive processes as reading and processing the musical note – the “language” specialized to music, as a result of their musical training and experience. In this sense, the Wernicke’s area may reflect musicians’ conscious or subconscious comprehension of the music note when hearing or imagining music.

Our results suggest that musical imagery and perception may also activate the Broca’s area, despite to a less degree than the Wernicke’s area. Although the Broca’s area is related to speech or language production^42,43^, recent findings also suggest its involvement in musical perception, especially for processing the musical syntax and structure ^44–46^. Instrumental music and language both have syntactic systems that use complex and hierarchically structured sequences and terminology. The processing of musical syntax and language syntax may share a common functional module at the Broca’s area^47^.

Our results also suggest that both musical imagery and perception evoke bilaterally symmetric activations and networks. This finding contradicts the earliest view that brain function related to music is mostly or exclusively specialized to the right hemisphere. Such a functional lateralization may depends on musical experiences ^48,49^. Studies have shown that extensive musical training shapes brain development ^50^, improves verbal intelligence ^51^, and enhances executive function ^52^. We speculate that the bilateral activation and network with musical imagery and perception indicate the breakdown of lateralization of music processing, as a result of experience-dependent neural plasticity.

The response at the Wernicke’s area was highly correlated with the auditory feature of the music as being imagined or heard (Fig. 3). This observation offers insights into the function of the Wernicke’s area: it encodes music information albeit whether it is internally generated (imagery) or externally stimulated (perception) in musicians, in addition to its known function in language comprehension in non-musicians^38,39^. In this study, the music information was expressed in terms of the spectral flux of the sound wave, indicating the dynamic changes of power spectrum. As such, this feature was auditory in nature, not directly conveyed by the music visualization. Thus, this feature representation was most likely generated by cortical processes underlying imagery rather than visual perception.

The Wernicke’s area also showed a response in correlation with the spectral flux when the music was directly heard. However, similar correlations were consistently observed beyond the Wernicke’s area, including the primary and secondary auditory cortex. The primary auditory cortex did not show any response correlated to this auditory feature during musical imagery (Fig. 3). This difference suggests that the primary auditory cortex encodes the auditory feature of a music, only when it is perceived but not when it is imagined.

Although the Wernicke’s area encoded auditory information in both musical imagery and perception, its functional connectivity appeared largely different in these two conditions (Fig. 4). For the musical perception task, the Wernicke’s area interacted with bilateral auditory cortices, showing a pattern of task-evoked functional connectivity consistent with intrinsic functional network in the resting state. However, during musical imagery, the Wernicke’s area engaged itself in a more complex pattern of functional connectivity, including a subset of the auditory network, the attention network, and the default-mode network.

This finding makes intuitive sense, as musical perception is a more natural and less demanding than musical imagery. The attention demand during musical imagery explains the positive interaction between the Wernicke’s area and the attention network ^53^. It also explains the negative interaction with the default-mode network, which is often deactivated by attention demanding tasks ^54^ and anti-correlated with the attention network at rest ^55^. The auditory network evoked by musical imagery involved the Wernicke’s area and the secondary auditory area, but skipped the primary auditory cortex, unlike the intrinsic auditory network or the task-evoked auditory network during musical perception. The absent of primary auditory cortex in musical imagery is consistent with most of the previous studies^20,24,56^, but not Kraemer et al., 2005^57^, which remains a topic under debate.

These findings provide a convincing demonstration that task-evoked functional networks may differ from intrinsic functional networks in the resting state, despite a general correspondence between them ^58^. A task-evoked network may break up an intrinsic network to involve a subset of regions, or may recruit multiple intrinsic networks through network-network interactions. The degree to which a task-evoked network deviates from its intrinsic mode might depend on the cognitive load and the frequency of occurrence of the task.

## Methods and Materials

### Subjects and Stimuli

Three healthy volunteers participated in the study with informed written consent obtained from every subject according to a research protocol approved by the Institutional Review Board at Purdue University. Every subject had many years of sustained musical training (Table1).

**Table1.**
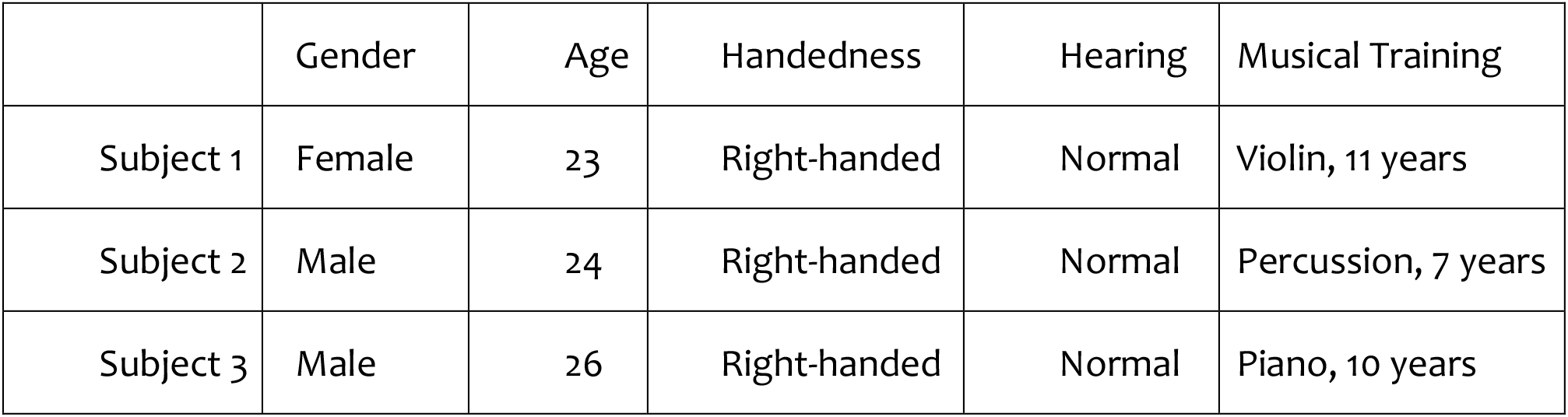
Subject information.

The auditory stimulus was the first 8 minutes from the first movement of Beethoven’s Symphony 9 (sampling rate: 11025 Hz), delivered through binaural MR-compatible headphones (Silent Scan Audio Systems, Avotec, Stuart, FL). This music piece was visualized as a movie through Stephen Malinowski’s Music Animation Machine ^59^. The visualization is a graphic music score based on the design of a 2-dimension piano roll. As illustrated in Fig. 1a, it is an animation with rectangular bars moving from right to left as the music flows; the current note is the highlighted bar in the middle of the screen. Every subject imagined the music while watching the movie that visualized the music. The movie offered the real-time visual cue to control the timing and inform the content of musical imagery. Prior to fMRI, each subject was trained for 1 to 2 hours, to be familiar with the music visualization and the imagery task. For this purpose, 1 to 2 hours of training was sufficient, as this visualization was intuitive for musicians as well as non-musicians ^60^. The visualized music was delivered through a binocular goggle system (NordicNeuroLab Visual System, Bergen, Norway) using Psychophysics Toolbox 3 (http://psychtoolbox.org).

For musical perception, each subject was instructed to listen to this music with attention; the same perception task was repeated 8 times. For musical imagery, each subject was instructed to imagine the music piece while watching the music visualization; the same imagery task was repeated 12 times.

### MRI Acquisition and Preprocessing

MRI data were acquired in a 3T MRI system (Signa HDx, General Electric Healthcare, Milwaukee) with a 16-channel receive-only phase-array surface coil (NOVA Medical, Wilmington). Structural MRI with T1 and T2-weighted contrast were both acquired with 1mm isotropic resolution. Functional MRI data were acquired by using a single-shot, gradient-recalled echo-planar imaging sequence with 3.5 mm isotropic spatial resolution and 2 second temporal resolution (38 interleaved axial slices with 3.5 mm thickness and 3.5×3.5 mm^2^ in-plane resolution, TR/TE=2000/35ms, flip angle=78°, field of view=22×22 cm^2^). For each subject was scanned with fMRI for 26 sessions: 8 with the musical perception task, 12 with the visually-cued musical imagery task, and 6 in the wakeful eyes-closed resting state. Every session lasted 8 minutes.

MRI/fMRI images were preprocessed using a similar pipeline as in the Human Connectome Project ^61^. Briefly, all fMRI images were corrected for slice timing and motion, aligned to structural images, normalized to the Montreal Neurological Institute (MNI) space, transformed onto individual cortical surfaces, and then co-registered based on myelin density and cortical folding patterns. After that, slow trend in the fMRI time series was corrected by regressing out a fourth-order polynomial function. Then fMRI data were temporally standardized (with a zero mean and unitary variance) and spatially smoothed by applying a 2-D Gaussian kernel of 2mm full width at half maximum.

### Mapping cortical activations during musical perception or imagery

For each subject, we mapped the cortical activation during musical perception or imagery. For either task, intra-subject reproducibility in voxel time series was calculated, as previously used elsewhere ^31,32^. This reproducibility was measured separately for each voxel as the temporal correlation between different sessions of the same task: 28 inter-session pairs among 8 sessions for the perception task, 66 inter-session pairs among 12 sessions for the imagery task. The significance of intra-subject reproducibility was evaluated by using a parametric statistical test based on linear mixed-effects (LME) modeling ^62^ implemented in AFNI (http://afni.nimh.nih.gov), a rigorous approach that accounts for the dependences among different inter-session pairs in the significance test. Specifically, the Fisher-transformed inter-session correlation *z_ij_* is expressed in a crossed random-effects model,

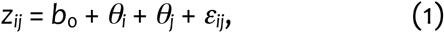

where *b*_0_ is the effect of interest, the average correlation across all inter-session pairs, *θ_i_* and *θ_j_* are the random effects associated with the *i*th and *j*th sessions, respectively, *ε_ij_* is the residual term, and *i* = 1, 2, …, *n* (*n* is 8 for perception and 12 for imagery).

### Mapping the common activations between musical perception and imagery

For each subject, we also mapped the cortical regions, where the musical perception and imagery tasks evoked the same response. For this purpose, we calculated for each voxel the temporal correlation in the voxel time series between every perception session and every imagery session. As such, the cross-task correlation was measured for a total of 96 pairs of sessions given 8 perception sessions and 12 imagery sessions. The significance of cross-task correlation was evaluated for each subject, based on a crossed random-effects LME model ^62^,

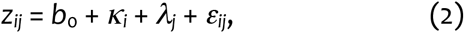

where *κ_i_* and *λ_j_* are the random effects are associated with the *i*th session of perception and the *j*th session of imagery, respectively, *i* = 1, 2, …, 8, and *j* =1, 2, …, 12.

### Mapping the fMRI correlates to auditory feature time series

We further explored the musical content coded in the fMRI response. For this purpose, the spectral flux of the music stimulus (in its auditory form) was extracted by the MIRtoolbox in Matlab ^44^. Spectral flux measures the change in power spectrum of a signal, which is calculated as the 2-norm (also known as the Euclidean distance) between the normalized spectra from adjacent frames^33^. The feature frame size was set to 1 second with a step size of 0.1 second. The spectral flux time series was convolved with the canonical hemodynamic response function (HRF) ^63^. The resulting time series was correlated with the fMRI signal at every voxel. For either the musical imagery or perception task, the correlation was calculated with the voxel time series averaged across all the repeated sessions of the same task.

### Mapping task-evoked networks during musical perception and imagery

For each subject, we mapped the task-evoked patterns of functional connectivity during either the musical perception or imagery task based on a seed-based inter-session correlation analysis ^34^. Specifically, the time series at a seed location was taken from one session; the seed time series was correlated with every voxel time series during a different session of the same task. Four seed locations were chosen: the left/right Wernicke’s area, and the left/right secondary auditory cortex (or the “belt” area). Such choices were based on the regions of interest (ROI) for both musical perception and imagery, as highlighted in our results (Fig. 1). The inter-session functional connectivity was evaluated for every pair of distinct sessions of the same task. The significance of inter-session functional connectivity was also evaluated using the LME method ^62^. See Eq. (1).

For each subject, we further compared the task-evoked networks with intrinsic functional networks observed with resting-state fMRI. Seed-based correlations in spontaneous activity were mapped with the same seed locations as mentioned above. The voxel time series was concatenated across the six resting state sessions for each subject, and temporal correlations were calculated based on the concatenated data. The statistical significance was evaluated based on two-tailed one sample t-test (corrected at false discovery rate (FDR) q<0.01).

## Data Availability

The datasets generated during and/or analyzed during the current study are available from the corresponding author on reasonable request.

## Acknowledgement

This work was supported in part by NIH R01MH104402 (PI: Liu) and by Purdue University. G. Chen was supported by the intramural program of National Institute of Mental Health.

## Authors’ Contributions

Y. Zhang and Z. Liu designed the study. Y. Zhang, H. Wen and K. Lu collected the data. Y. Zhang analyzed the data. Y. Zhang and G. Chen developed the analysis methods. Y. Zhang, Z. Liu, and G. Chen wrote the paper. All authors contributed to data interpretation.

